# Optimising analysis choices for multivariate decoding: creating pseudotrials using trial averaging and resampling

**DOI:** 10.1101/2023.10.04.560678

**Authors:** C. L. Scrivener, T. Grootswagers, A. Woolgar

**Affiliations:** MRC Cognition and Brain Sciences Unit, University of Cambridge, Cambridge, UK; School of Psychology and Neuroscience, University of Glasgow, Glasgow, UK; The MARCS Institute for Brain, Behaviour and Development, Western Sydney University, NSW, Australia; School of Computer, Data and Mathematical Sciences, Western Sydney University, NSW, Australia; Department of Psychology, University of Cambridge, UK

**Keywords:** Multivariate pattern analysis, decoding, pseudotrials

## Abstract

Multivariate pattern analysis (MVPA) is a popular technique that can distinguish between condition-specific patterns of activation. Applied to neuroimaging data, MVPA decoding for inference uses above chance decoding to identify statistically reliable condition-specific information in neuroimaging data which may be missed by univariate methods. However, several analysis choices influence decoding results, and the combined effects of these choices have not been fully evaluated. In particular, an increasingly popular approach is to average data from several trials together before training an MVPA classifier, but the decision about how much averaging to do is arbitrary and the effect of varying this parameter has not been documented. Here we systematically assessed the influence of trial averaging and resampling on decoding accuracy and subsequent statistical outcome on simulated data. Although the optimal parameters varied with the classifier and cross-validation approach used, we found that modest trial averaging using up to 5-10% of the total number of trials per condition improved decoding accuracy and associated t-statistics. In addition, a small amount of resampling could improve t-statistics and classification performance, but was not always necessary. We provide code to allow researchers to optimise these analysis choices for the parameters of their data.

## Introduction

The last decade has seen an explosion in the popularity of multivariate pattern analysis (MVPA) for neuroimaging data. By identifying condition-specific patterns of activation, MVPA can reveal the evolution of information processing over time and/or space and is sensitive to information missed by univariate methods (e.g., Grootswagers, 2017; Pereira et al., 2009; Haynes & Rees, 2006). Typically, cognitive neuroscience experiments employ MVPA techniques, such as linear classification (“decoding”), to make inferences about the type of information decodable from neuroimaging data, and to characterise these “neural representations” in terms of when and where they can be decoded, whether they generalise between conditions, and if they change with experimental manipulations. Decoding for inference typically compares decoding metrics (e.g., classification accuracy) between conditions or to chance, drawing inference about neural processing from statistically reliable condition-specific information in neuroimaging data.

There are many ways to run a decoding for inference analysis, and multiple decisions are likely to influence decoding success. One analysis option is the creation of ‘pseudotrials’, or ‘supertrials’, by averaging data from subsets of trials together before performing MVPA. Previous results demonstrate that pseudotrial averaging can lead to an increase in classification accuracy, compared to single-trial decoding (e.g., Adam, 2020; Tuckute, 2019; Hebart et al, 2018; Grootswagers, 2017; Isik, 2014). However, too much averaging can be detrimental, as increasing the number of trials per pseudotrial can also increase the between-subject variance in decoding accuracy, which in turn affects the statistical outcome. A second decision is the cross-validation procedure used to evaluate the generalisation of classification across subsets of the data (Bishop & Nasrabadi, 2006). In a leave-one-trial-out procedure, the number of cross-validation folds is equal to the number of trials available per exemplar. At each fold, the classifier is trained on all but one of the trials, which is then used to test the classification. A leave-one-pseudotrial-out method uses the same logic, but is based on averaged subsets of the original trials. Another option is to divide the data into larger chunks or blocks across which to train and test the classifier. For example, in a study with 90 trials per exemplar, 10-fold cross-validation would split the data into 9 sets of 10 trials to use iteratively for training and testing. This means that at each iteration, the classifier is trained on fewer data points than for leave-one-trial-out, but previous work has shown that this may provide more stable estimates with lower variance across decoding accuracies (Varoquaux et al., 2017). When combined with trial averaging, pseudotrials can be created either by averaging all of the trials within a chunk or by grouping them into smaller subsets of trials. Therefore, at least two interacting parameters seem likely to affect results: the number of trials averaged together before classification, and the number of folds into which the data are split for cross-validation.

Thus far the choice of these parameters has been largely arbitrary, resulting in a wide variety of trial averaging and cross-validation approaches in the literature. To name a few examples, in Bae and Luck (2018) and Foster et al. (2017a), the available trials per condition were randomly divided into 3 chunks before being averaged and decoded using a 3-fold cross-validation. In Isik et al. (2014), groups of 10 trials were averaged before using a 5-fold cross-validation, and in Duncan et al. (2023), groups of 3 trials were averaged before using a 10-fold cross-validation. Goddard et al. (2022) averaged over 16 trials, before using an 8-fold cross-validation, and Petit et al. (2023) used the median of 5 trials and a leave-one-pseudotrial-out cross-validation with 22 folds. Given the variation across experiments, it can be difficult to make decoding analysis decisions regarding a new set of data.

In an evaluation of different decoding approaches, Grootswagers et al. (2017) compared decoding accuracies when averaging together 4, 8, 16, and 32 trials (equivalent to creating 8, 4, 2, and 1 pseudotrials) from an experiment of 32 trials per condition (64 total trials). A leave-one-pseudotrial-out cross-validation scheme was used, meaning that the number of cross-validation folds was determined by the number of pseudotrials. All averaging procedures increased decoding accuracies compared to no trial averaging, but the least amount of averaging (4 trials, 12.5%) provided the best trade-off between signal-to-noise and number of pseudotrials. In addition, they found similar decoding accuracies for 10-fold and leave-one-trial-out cross-validation methods. They reported lower decoding accuracies using a 2-fold cross-validation procedure, though others have argued that this approach may be associated with higher statistical power overall (Valente et al., 2021). In a different implementation, Adam et al. (2020) compared trial averaging results using a 3-fold classification that was independent from the number of pseudotrials created. They averaged groups of trials within each fold, ranging from 5 to 25, and found a significant increase in the average decoding accuracy with more averaging. However, given the associated increase in between-subject variance, they concluded that the average of 10 trials (this corresponded to between 2.5 and 10% of the available trials across datasets of different sizes) was optimal for the number of trials available. Thus, while there is clearly a trade-off between providing a classifier with fewer less noisy trials (more averaging) or more noisy trials (less averaging), it is not yet clear how to optimise this decision. In addition, it is unclear whether and how the optimal amount of averaging depends on other factors such as number of trials, choice of classifier, effect size, etc.

The aim of the current work was to inform future decoding studies by systematically assessing the influence of a range of parameters on decoding accuracy and subsequent statistical outcome, using data simulated in CosMoMVPA (Oosterhof, 2016). We varied several parameters across simulations, including the number of trials per pseudotrial, cross-validation procedure, number of trials per condition and the size of the underlying effect, and assessed the data for three different linear classifiers. Decoding results varied with the classifier used, the cross-validation approach, and the number of the trials averaged to create each pseudotrial. In addition, we evaluated the influence of random sampling with replacement (‘resampling’) on decoding accuracy, which has not yet been comprehensively assessed. We hypothesised that this would be beneficial when creating pseudotrials as it increases the number of samples available for classification, which would normally be diminished by trial averaging. Although the optimal parameters varied with classifier and cross-validation approach, we found that using roughly 5-10% of the total number of trials per condition was optimal for creating pseudotrials. In addition, a resampling value of 2 could improve t-statistics and reduce the impact of the number of trials per pseudotrial on classification performance.

## Methods

We used CoSMoMVPA (Oosterhof, 2016) to simulate multiple datasets, each with two experimental conditions, and examined the influence of a) the number of trials used per pseudotrial, b) the number of times each trial was resampled, c) the simulated class distance, and d) the choice of classifier. We chose three popular classifiers that are implemented in CoSMoMVPA: linear support vector machine (libSVM), linear discriminant analysis (LDA), and Gaussian Naïve Bayes. For experiment 1, we simulated data with 700 features (reflecting channels/voxels) from 100 ‘subjects’ with 2 conditions and 90 trials per condition. To check whether the results were dependent on the specific number of trials per condition, we also simulated a second experiment with only 45 trials per condition. We manipulated the multivariate class distance using built-in CoSMoMVPA functions: individual values were drawn from a normal distribution (sd=1) with the amount of separation between class-means for each feature defined as *class-mean = class_distance/sqrt(log(700*ntrials))* with *ntrials* either 90 or 45 (for the 2 experiments), and used 3 values for *class_distance* (0, 0.1, and 0.2). We refer to a simulated class distance of 0 as ‘no effect’, 0.1 as a ‘small effect’, and 0.2 as a ‘large effect’. Results for a smaller number of participants (50) and fewer features (6) can be found in the supplementary material (S4 and S5). Simulation scripts are available on the Open Science Framework, https://osf.io/hjf75/.

We ran both a ‘chunking’ and a ‘leave-n-pseudotrials-out’ cross-validation procedure as both approaches are commonly used. For the ‘chunking’ procedure, we created pseudotrials separately within each of 3 ‘blocks’ of trials, and used a 3-fold cross-validation method (additional results for a 10-fold cross-validation procedure can be found in the supplementary material S6). For the ‘leave-n-pseudotrials-out’ method, the number of cross-validation folds was determined by both the number of trials per pseudotrial and the amount of trial resampling (*number of folds = total trial number/trials per average*trial resampling*, with a fixed minimum of 2 folds). The more trials averaged together, the fewer the total number of pseudotrials, and therefore the fewer possible folds. The more trial resampling, the more trials were allocated to each fold to prevent all pseudotrials from being identical, and therefore the fewer possible folds. The number of pseudotrials included in each cross-validation fold, or ‘n’ pseudotrials, was determined by the number of times each trial was resampled (the more times each trial was resampled, the more pseudotrials could be created per fold).

In experiment 1 (90 trials/condition), for both methods we resampled each trial between 1 and 15 times and used between 1 and 30 of the available trials per pseudotrial. In Experiment 1 (45 trials/condition), we resampled each trial between 1 and 10 times and used between 1 and 15 of the available trials per pseudotrial. Any trials that could not be used in particular iteration of pseudotrial creation (this happened when the number of trials was not perfectly divisible by the number of trials per pseudotrial) were left out of that iteration of the analysis. To ensure that the results were not dependent on the specific division of trials into pseudotrials (Goddard et al., 2022), for each ‘subject’ and parameter set, we ran 100 iterations of the pseudotrial procedure and averaged the 100 resulting classification accuracies to give a single value for that subject and parameter set. This repetitive iteration of pseudotrial creation reduced the additional variance introduced by the averaging procedure and therefore appears to be essential to derive benefit from the pseudotrial scheme (see Supplementary Figure 8 where we present results using different numbers of iterations). For all simulations, we report the average decoding accuracy across subjects, the standard deviation, and their ratio (t-score against chance level decoding across subjects, 50%).

## Results

### Influence of trial averaging

First, we considered the influence of trial averaging. We used a 3-fold cross-validation procedure, in which trials were randomly allocated to one of three blocks before pseudotrials were created. Every analysis used the same 3-fold cross validation procedure, and only the number of pseudotrials per block varied across averaging and resampling values. Importantly, for the dataset with no simulated difference between conditions (**Figure 1A**) average decoding accuracy remained at 50% with the creation of pseudotrials. This reassures us that averaging the data cannot create effects where they are not present in the underlying data.

**Figure 1.**
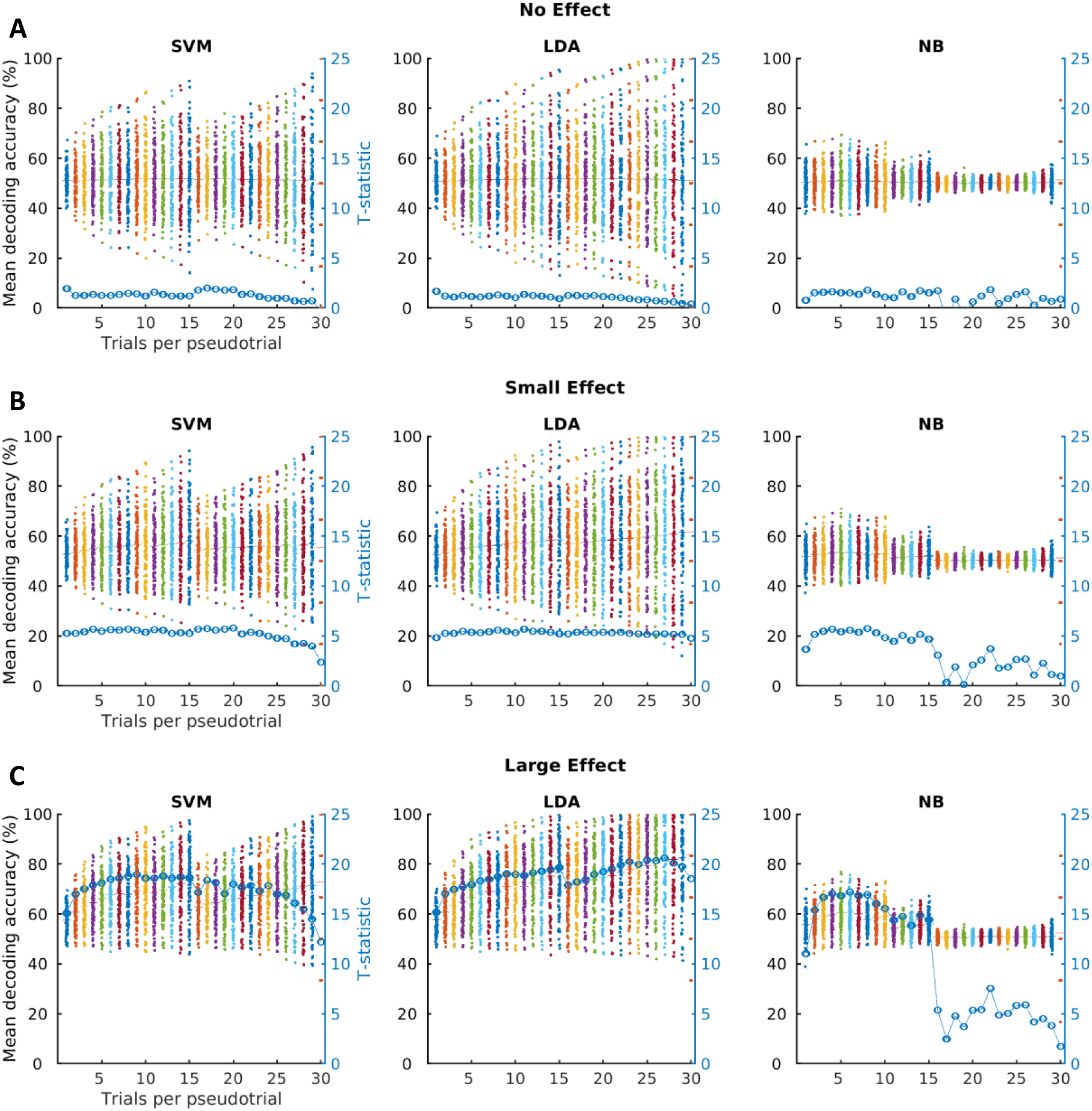
The influence of trial averaging, size of simulated effect, and classifier type on decoding accuracy and t-statistics. Rows correspond to the three simulated effects; **A**) no effect, **B**) small effect, and **C**) large effect. Columns correspond to results from the three classifiers tested (SVM = support vector machine, LDA = linear discriminant analysis, NB = Naïve Bayes). We simulated data from 100 subjects with 90 trials per condition. Pseudotrials were created separately within 3 ‘blocks’ of trials, facilitating a 3-fold cross-validation approach, with trials randomly allocated to pseudotrials across 100 iterations of pseudotrial creation for each subject. ‘Trials per pseudotrial’ indicates the number of trials used to create pseudotrials for each of the two simulated conditions, which was always matched. Using one trial per pseudotrial (the first value on the x-axis) is equivalent to no trial averaging and using all 30 individual trials per condition per ‘block’. Each datapoint represents the average cross-validated decoding accuracy for one ‘subject’. The t-statistic (blue line and scale bar) is to quantify the trade-off between modulating the mean decoding accuracy and variance over ‘subjects’. It was calculated at the group-level by subtracting 50 (theoretical chance level) and dividing by the standard error.

For the data with small and large differences between conditions (**Figure 1B**, **Figure 1C**), we found that averaging even a few trials together was beneficial for classification performance, resulting in increased decoding accuracy and higher t-statistics, despite the concurrent increase in standard deviation. However, optimising this was non-trivial. Instead, there was a complex pattern of results that varied with the number of trials per pseudotrial, classifier type, and size of effect. Including too many of the available trials in an average (reducing the number of datapoints available to the classifier) could be detrimental, as the increase in standard deviation outstripped the increase in average decoding, resulting in reduced t-values. Using more than half of the available trials per pseudotrial was particularly detrimental for the Naïve Bayes classifier, which benefited from less averaging and more samples for the classifier. This was also the case for the SVM classifier, but to a lesser extent, and most visible when the class distance was the greatest.

### Influence of trial resampling

Next, we examined the influence of trial resampling and its interaction with the number of trials per pseudotrial. Once again, with no simulated effect, decoding remained at 50%, reassuring us that averaging and resampling the data cannot create effects where there are none (Supplementary Figure 1). For the data with a small simulated difference between conditions (Figure 2), we found that using up to a third of the original trials per chunk to create each pseudotrial was sufficient to aid decoding (i.e., 10 of the 30 trials per chunk, or 11% of the total 90 trials). Combining this with a small amount of resampling further increased decoding accuracy, presumably because resampling allows more pseudotrials to be created. For example, with 30 trials per condition in each of the 3 chunks, averaging together 10 trials creates 3 pseudotrials per chunk if no resampling is used. With a resampling value of 2, each trial is included in 2 pseudotrials, meaning that 6 pseudotrials are created per chunk. While a resampling of 2 increased t-values for the Naïve Bayes classifier, this was not the case for SVM and LDA, where resampling mostly acted to stabilise the influence of the number of trials per pseudotrial. The results for a large effect were similar and can be found in Supplementary Figure 2.

**Figure 2.**
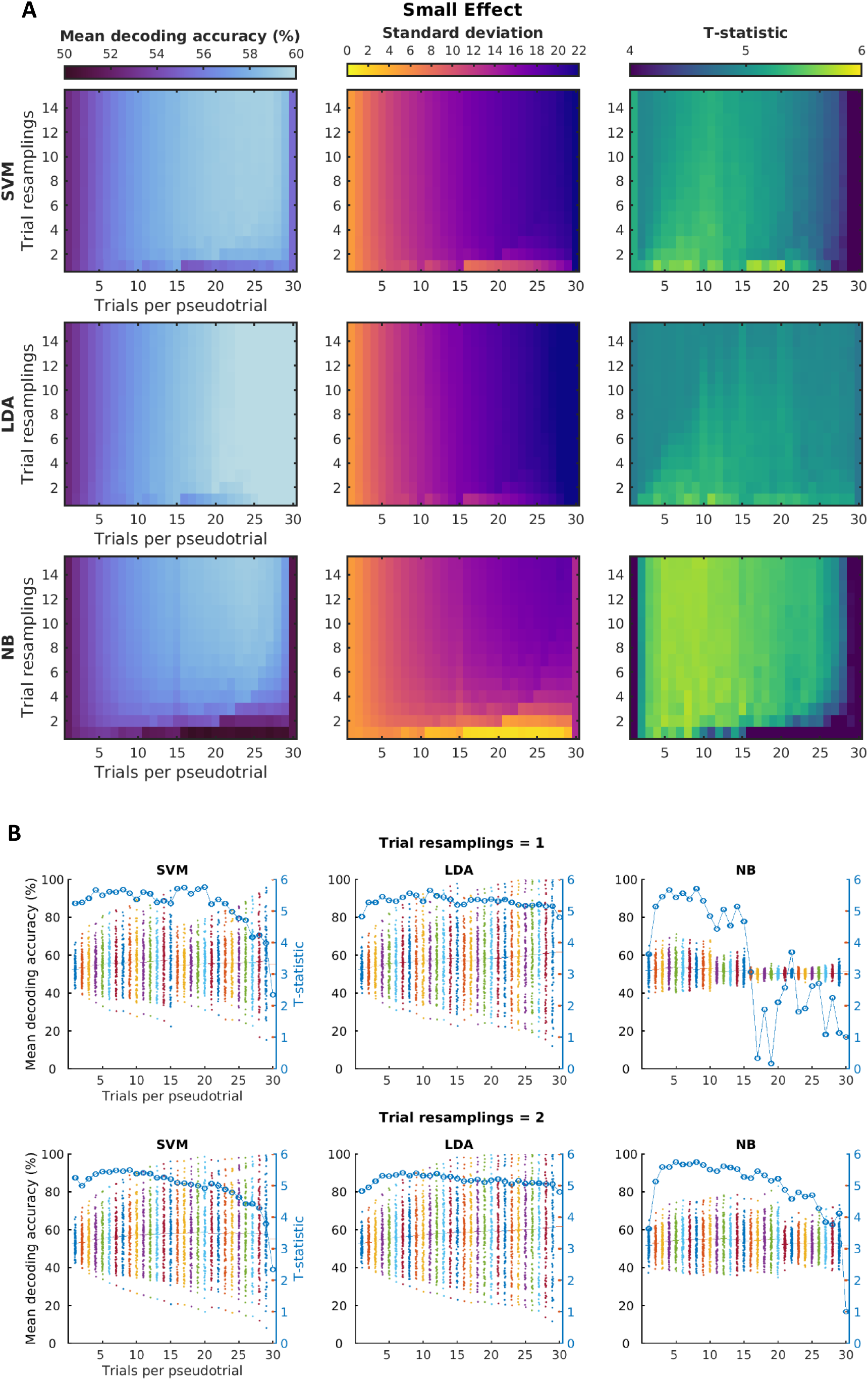
**A**) The influence of averaging and resampling on decoding accuracy (left column), standard deviation across subjects (middle column), and group-level t-statistics (right column). Each cell represents the outcome across subjects for one combination of trial averaging and resampling values. Lighter colours on all three metrics correspond to a better decoding outcome (i.e. higher decoding values, lower standard deviation, and higher t-statistics). Rows correspond to results from the three classifiers tested (SVM = support vector machine, LDA = linear discriminant analysis, NB = Naïve Bayes). Pseudotrials were created separately within 3 allocated ‘blocks’ of trials, facilitating a 3-fold cross-validation approach, with trials randomly allocated across 100 iterations of pseudotrial creation. For the results plotted here, we simulated data from 100 subjects with 90 trials per condition and a small effect between condition (see supplementary Figures 1a and 1b for the results with no effect and a large effect). **B**) To further illustrate the influence of trial resampling value on decoding outcomes, we also show the scatter plots for resampling values of 1 (top row, reproduced from Figure 1B) and 2 (bottom row). These correspond to the data in the bottom two rows of the plots shown in 2A. Each datapoint represents the cross-validated decoding accuracy for one subject. The t-statistic was calculated at the group-level to test the decoding accuracy against chance across all subjects (50%).

### Influence of fewer trials per condition

Next, we checked whether the same principles would apply to an experiment with fewer trials per condition. Figure 3 demonstrates the effect of trial resampling on a smaller dataset with only 45 trials per condition, meaning 15 trials in each of the 3 chunks. Once again, averaging up to a third of the original trials per chunk (i.e., 5 trials or less) to create each pseudotrial was sufficient to aid decoding performance. A small amount of resampling combined with trial averaging aided the classification particularly for the Naïve Bayes classifier (Figure 3B). However, high values on either parameter was generally detrimental for classifier performance; although the decoding accuracies were raised, an increase in standard deviation resulted in a decrease in t-values (Figure 3A**, column 3**).

**Figure 3.**
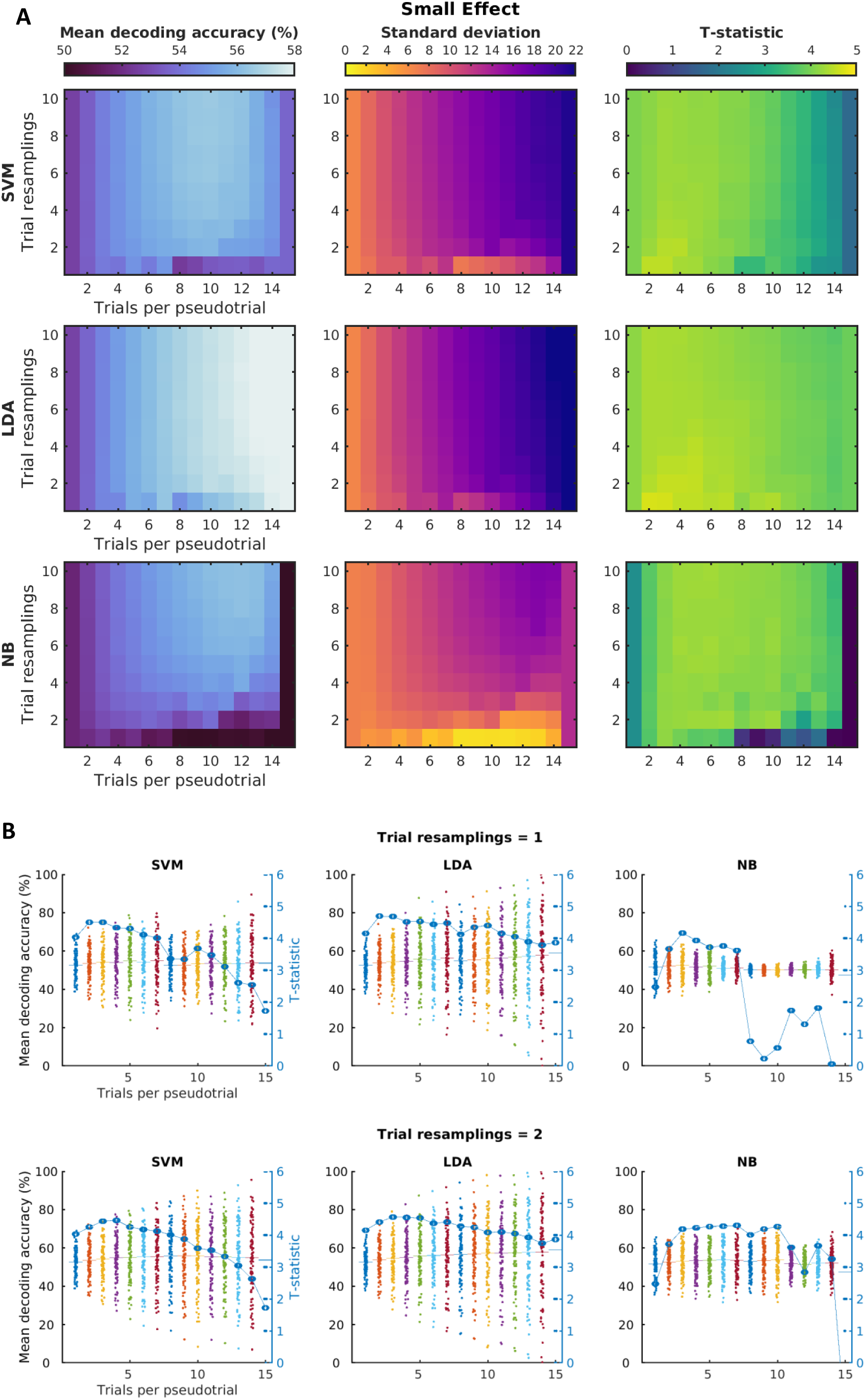
**A**) The influence of averaging and resampling on decoding accuracy (left column), standard deviation across subjects (middle column), and group-level t-statistics (right column). Each cell represents the outcome across subjects for one combination of trial averaging and resampling values. Lighter colours on all three metrics correspond to a better decoding outcome (i.e. higher decoding values, lower standard deviation, and higher t-statistics). Rows correspond to results from the three classifiers tested (SVM = support vector machine, LDA = linear discriminant analysis, NB = Naïve Bayes). Pseudotrials were created separately within 3 allocated ‘blocks’ of trials, facilitating a 3-fold cross-validation approach, with trials randomly allocated across 100 iterations of pseudotrial creation. For the results plotted here, we simulated data from 100 subjects with 45 trials per condition and a small effect (see Supplementary Figure 2 for a large effect). **B**) To further illustrate the influence of trial resampling value on decoding outcomes, we also show the scatter plots for resampling values of 1 and 2. These correspond to the data in rows 1 and 2 of the plots shown in A. Each datapoint represents the cross-validated decoding accuracy for one subject. The t-statistic was calculated at the group-level to test the decoding accuracy against chance across all subjects (50%).

### Influence of a ‘leave-n-pseudotrials-out’ decoding approach

Next, we examined the alternative ‘leave-n-pseudotrials-out’ procedure, where the number of folds was determined by the number of pseudotrials created (number of folds = total trial number/(trials per average*trial resampling)). As shown in Figure 4, the largest t-statistics were found across all classifiers when using roughly 5% of the total 90 trials per pseudotrial (i.e., 4 or 5 trials) with a resampling of 2. Therefore, fewer trials per average were necessary to aid classification in the ‘leave-n-pseudotrials-out’ approach, compared to the optimal 11% for the 3-chunk version. The maximum decoding accuracies and t-values achieved using the ‘n-pseudotrials-out’ method were also slightly higher overall than the ‘3-block’ procedure, but had more variation across the parameter space. This is presumably due to the higher number of folds that could be used at low values of averaging and resampling. However, once the number of folds reached the minimum of 2, the influence of the parameters was reduced. This occurred when the combined averaging and resampling parameters exceed the number of trials that could be allocated to more than 2 folds.

**Figure 4.**
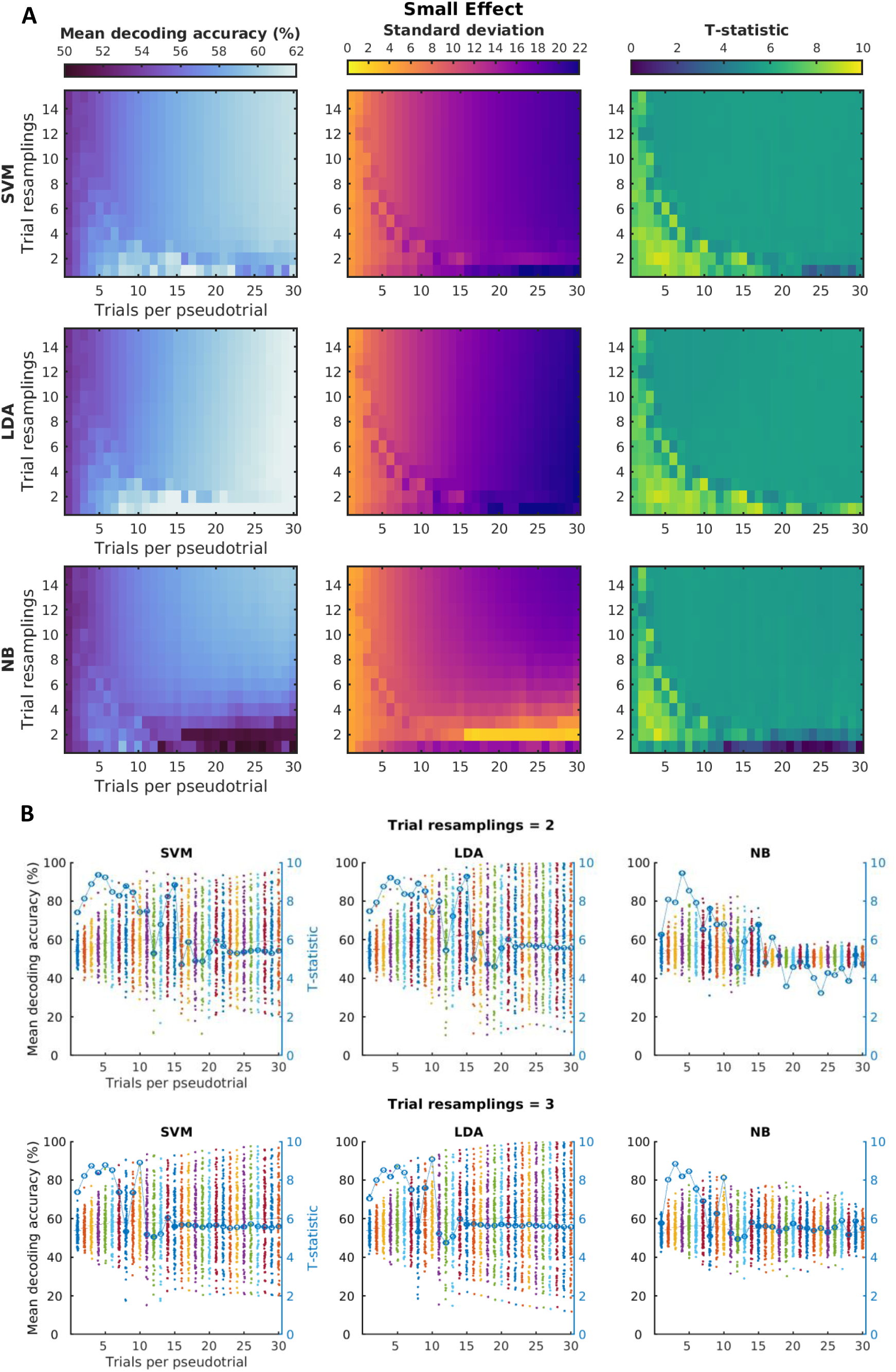
**A**) The influence of averaging and resampling using a ‘leave-n-pseudotrials-out’ approach on decoding accuracy (left column), standard deviation across subjects (middle column), and group-level t-statistics (right column). Each cell represents the outcome across subjects for one combination of trial averaging and resampling values. Lighter colours on all three metrics correspond to a better decoding outcome (i.e. higher decoding values, lower standard deviation, and higher t-statistics). Rows correspond to results from the three classifiers tested (SVM = support vector machine, LDA = linear discriminant analysis, NB = Naïve Bayes). Here the number of cross-validation folds was determined by the number of trials per pseudotrial (the more trial averaging and resampling, the fewer possible folds), with trials randomly allocated across 100 iterations of pseudotrial creation. For the results plotted here, we simulated data from 100 subjects with 90 trials per condition and a small effect (see Supplementary Figure 3 for a large effect). **B**) To further illustrate the influence of trial resampling value on decoding outcomes, we also show the scatter plots for resampling values of 1 and 2. These correspond to the data in rows 1 and 2 of the plots shown in A. Each datapoint represents the cross-validated decoding accuracy for one subject. The t-statistic was calculated at the group-level to test the decoding accuracy against chance across all subjects (50%).

## Discussion

We examined the influence of averaging and resampling across several parameters, including classifier type, size of simulated effect, number of trials available per condition, and the cross-validation procedure. While increasing the number of trials used per pseudotrial generally increased decoding accuracy, it also increased the between-subject variance. We used a t-test to quantify the trade-off between these factors, and found that, with a ‘3-chunk’ or ‘10-chunk’ approach, using up to a third of the original number of trials per chunk (or ∼10% of all trials for 3-chunk, ∼3% of all trials for 10-chunk) aided decoding. This was consistent for both experiment sizes. For the 3-chunk procedure, there was an additional stabilising effect provided by a low resampling value of 2. This was equivalent to creating 6 pseudotrials within each of the 3 chunks (18 pseudotrials in total). However, too much averaging could be detrimental, particularly for the Naïve Bayes classifier, and little was gained by using high resampling values except to compensate for too much averaging using NB.

For the ‘leave-n-pseudotrials-out’ cross-validation approach, fewer trials were needed per pseudotrial to optimise the decoding outcome. Using around a sixth of the original number of trials per chunk (or 5% of all trials) produced the largest t-statistics across all classifiers, when combined with a low resampling value of 2. Higher decoding accuracies and t-statistics were achieved for some parameters using the ‘leave-n-pseudotrials-out’ approach compared to the ‘3-chunk’ version, but the different number of decoding folds resulted in more variation across the parameter space.

Although there were similarities across the three classifiers used, they responded differently across the parameter space, presumably due to the differences in their functions. Linear SVM separates classes by positioning a decision hyperplane in pattern space (Misaki et al., 2010). This hyperplane is chosen by maximising the distance to the patterns on either side, using the most informative data points that lie closest to the decision boundary (support vectors). Because of this, the SVM is not as influenced by changes in data points sitting away from the decision boundary. Therefore, SVM can perform well with limited data and will benefit most from having a few stable estimates near the decision boundary (Mur et al., 2009). Averaging may therefore aid classification using SVM by stabilising the data used to position the hyperplane.

In LDA, the pattern space is constructed by maximising the between-class variance while minimising the within-class variance. The hyperplane is positioned in the middle of the class means, assuming that the two classes have Gaussian distributions and equal covariance. Therefore, a change in any data point will shift the decision boundary and potentially influence the classification result. Gaussian Naïve Bayes is similar to LDA, but also assumes that there are no correlations between pairs of data points within the same class (zero off-diagonal covariance). Having a low number of data points in each class distribution may more negatively impact the performance of Gaussian Naïve Bayes (Misaki et al., 2010), as demonstrated here.

Here, we focused on t-scores derived from decoding accuracy as a quantification of the trade-off between increasing mean decoding and variance. Other metrics of class separation have been proposed, such as cross-validated Mahalanobis distance (Walther et al., 2016). We believe that our results would generalise to such other measures, as our results show that averaging increases separation between datapoints, evidenced by increased accuracy, which would similarly increase Mahalanobis distance. However, a full exploration of the effect of averaging on these other measures is outside the scope of the current paper. We also acknowledge that real neuroimaging data is more complex than the data simulated here, but aim to provide a starting point for researchers to examine the effects of decoding parameters in the simplest case.

In summary, we found that modest trial averaging can improve decoding accuracy and associated t-statistics, and that a small amount of resampling helps to stabilise the benefit of doing so. However, only a low resampling value is helpful, and is not always necessary. In addition, the use of pseudotrials did not increase decoding accuracies when no effect was present. Although we provide general guidelines, the optimal parameter choice (particularly, the number of trials per pseudotrial) will be data and design specific, so we provide analysis code for others to run simulations based on their own design and hypothesised effects. Our guidelines and code can be used to inform future multivariate brain decoding studies.

## Supplementary Material

Here we present additional results for effect sizes that were not included in the main text. This includes the influence of trial resampling, fewer trials per condition, and a leave-n-pseudotrials-out approach. We also present additional analyses that were not addressed in the main text. This includes the influence of a smaller number of subjects, a smaller number of features, increasing the number of cross-validation folds to 10, and using different iterations of random trial allocation to pseudotrials.

### S1. Influence of trial resampling (additional no effect and large effect)

Here we examined the influence of trial resampling on decoding accuracy when there was no simulated class difference. This differs from **main** Figure 2 which displayed the results for a small effect (class distance of 0.1). Decoding performed on data with no simulated effect remained at 50%, reassuring us that averaging and resampling the data cannot create effects where there is none (Supplementary Figure 1).

**Supplementary Figure 1.**
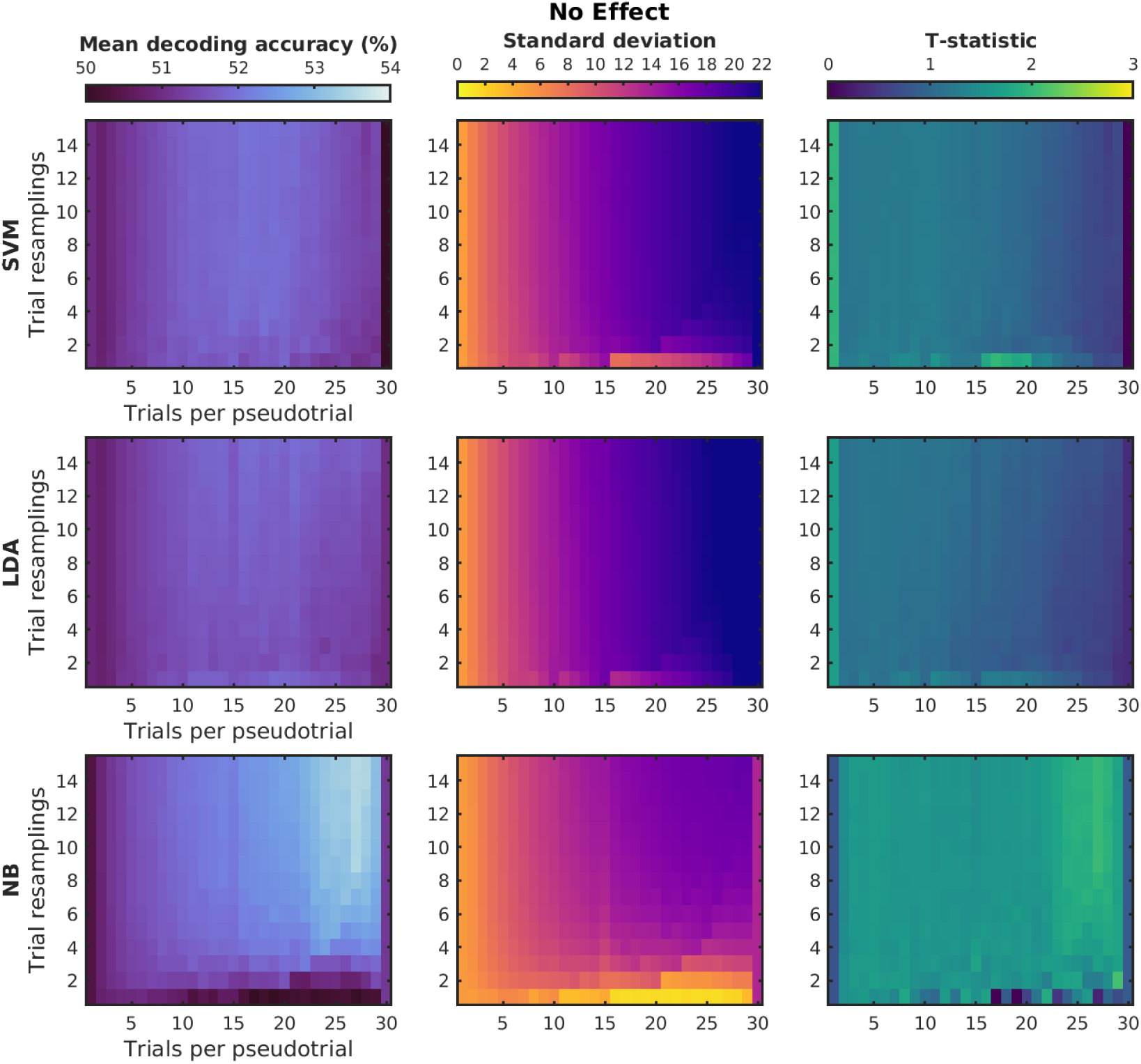
The influence of both averaging and resampling, on decoding accuracy, standard deviation, and t-statistics when the simulated data has no underlying effect. Rows correspond to results from the three classifiers tested (SVM = support vector machine, LDA = linear discriminant analysis, NB = Naïve Bayes). Pseudotrials were created separately within 3 allocated ‘blocks’ of trials, facilitating a 3-fold cross-validation approach, with trials randomly allocated across 100 iterations of pseudotrial creation. For the results plotted here, we simulated data from 100 subjects with 90 trials per condition with no effects, or a class distance of 0 (see main Figure 2 for a small effect, or class distance of 0.1).

For the data with a small simulated difference between conditions of 0.1 (**main** Figure 3) we found that using around a third of the original trials per chunk to create each pseudotrial (i.e., 10 trials) was sufficient to aid decoding. Combining this with a small amount of resampling further increased decoding accuracy. As shown in Supplementary Figure 2, this was also true for the data with a large effect (a class distance of 0.2), when using the Naïve Bayes classifier. These parameters also performed well for the SVM and LDA classifiers, but SVM performed equally with up to 50% of the original trials per pseudotrial (i.e., 15 trials) and a resampling value of 2. The LDA classifier performed well across a large range of parameters, providing that the resampling value was not higher than the number of trials per pseudotrial. With the larger simulated effect, the classifiers appeared to be more robust to the choice of averaging and resampling parameters.

**Supplementary Figure 2.**
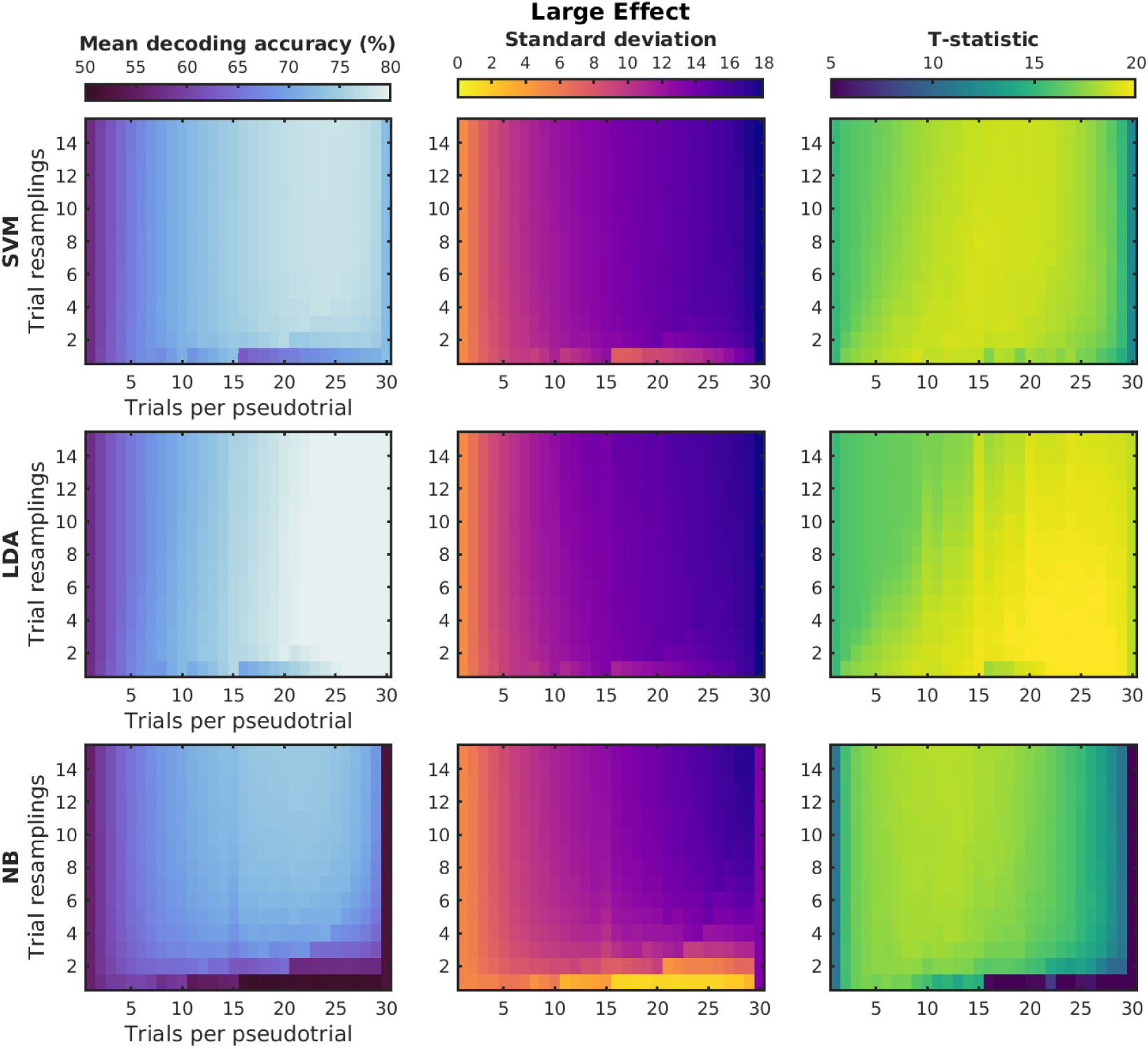
The influence of both averaging and resampling, on decoding accuracy, standard deviation, and t-statistics for simulated data with a large underlying effect. Rows correspond to results from the three classifiers tested (SVM = support vector machine, LDA = linear discriminant analysis). Pseudotrials were created separately within 3 allocated ‘blocks’ of trials, facilitating a 3-fold cross-validation approach, with trials randomly allocated across 100 iterations of pseudotrial creation. For the results plotted here, we simulated data from 100 subjects with 90 trials per condition and a large effect, or a class distance of 0.2 (see main Figure 2 for a small effect, or a class distance of 0.1).

### S2. Influence of fewer trials per condition (additional large effect)

Here we examined the effect of trial resampling on a smaller dataset with only 45 trials per condition, meaning 15 trials in each of the 3 chunks. This differs from **main** Figure 3 which displayed the results for small effect (a class distance of 0.1). Once again, a small amount of resampling combined with trial averaging aided the classification. High values on either parameter was detrimental for classifier performance, and using up to a third of the original trials (i.e., 5 trials or less) per pseudotrial with a resampling of 2 was optimal for both a small effect (**main** Figure 3), and a large effect (Supplementary Figure 3).

**Supplementary Figure 3.**
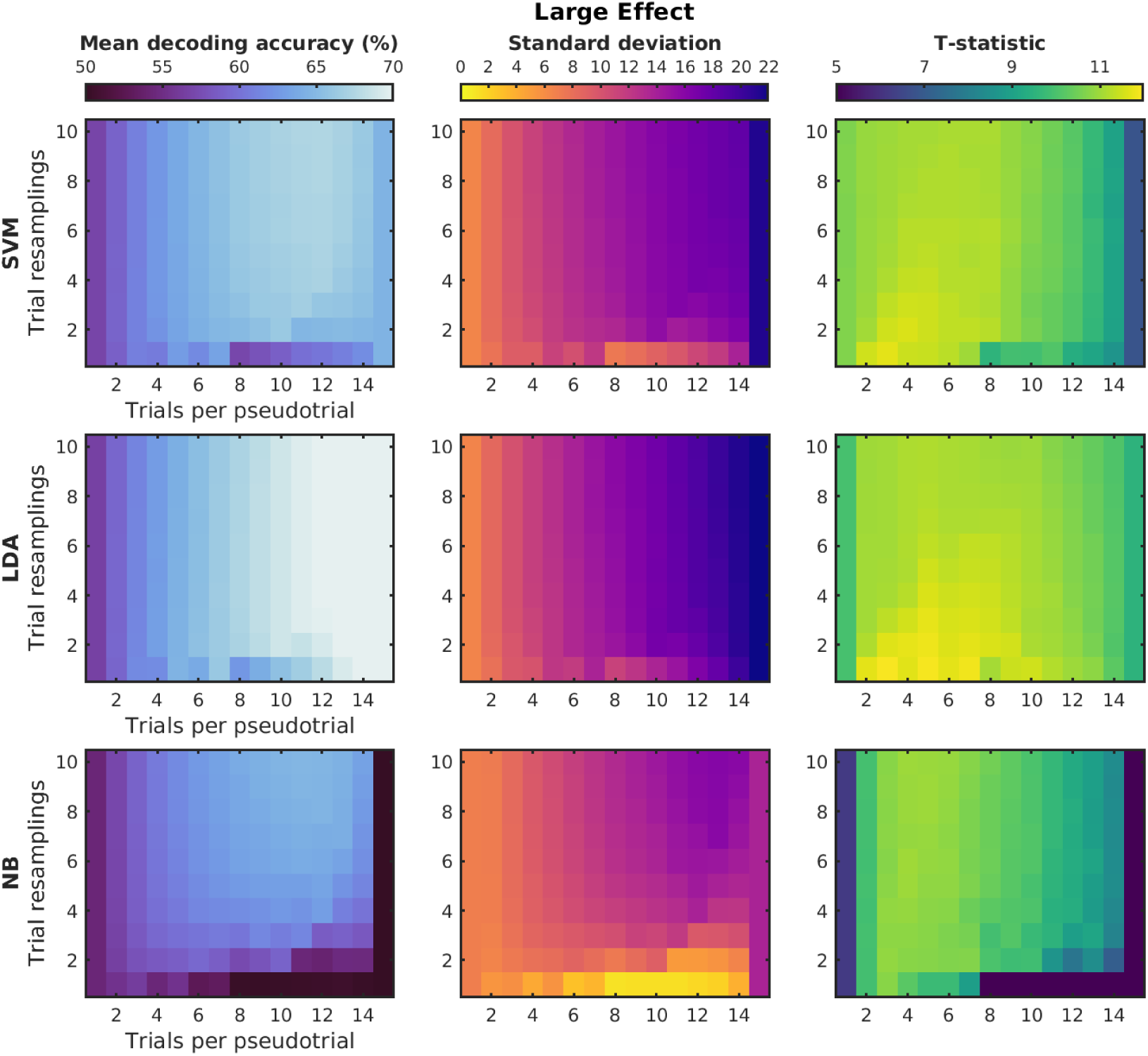
The influence of both averaging and resampling for a dataset with only 45 trials per condition for simulated data with a large underlying effect. Rows correspond to results from the three classifiers tested (SVM = support vector machine, LDA = linear discriminant analysis, NB = Naïve Bayes). Pseudotrials were created separately within 3 allocated ‘blocks’ of trials, facilitating a 3-fold cross-validation approach, with trials randomly allocated across 100 iterations of pseudotrial creation. For the results plotted here, we simulated data from 100 subjects with 45 trials per condition and large effect, or a class distance of 0.2 (see main Figure 3 for a small effect, or a class distance of 0.1).

### S3. Influence of a ‘leave-n-pseudotrials-out’ decoding approach (additional large effect)

Here we examined a ‘leave-n-pseudotrials-out’ procedure, where the number of folds was determined by the number of pseudotrials created, for a higher simulated class distance. This differs from **main** Figure 4 which displayed the results for small effect (class distance of 0.1). For the Naïve Bayes classifier, high t-statistics were achieved when using roughly 15% of the original trials per pseudotrial (i.e., 4 or 5 trials) with a resampling of 2 (Supplementary Figure 4). This is similar to the results with a small effect, or a class distance of 0.1 (**main** Figure 4). With a large effect (class distance of 0.2), the SVM classifier performed well with up to a third of the original trials (i.e., 10 trials) with a resampling of 2. For the LDA classifier, this could increase to half of the original trials (i.e., 15 trials) with a resampling of 2.

### S4. Influence of fewer ‘subjects’

Here we examined the influence of reducing the number of simulated ‘subjects’ from 100 to 50, which was not examined within the main text (Supplementary Figure 5). A similar pattern is found to the data in Supplementary Figure 2 with 100 subjects and a large effect (a class distance of 0.2), although the overall performance is reduced. For the Naïve Bayes classifier, we found that using around a third of the original trials per chunk to create each pseudotrial (i.e., 10 trials) was sufficient to aid decoding. Combining this with a low resampling value of 2 further increased decoding accuracy. The SVM and LDA classifiers performed well with up to 50% of the original trials per pseudotrial (i.e., 15 trials) and a resampling value of 2.

**Supplementary Figure 4.**
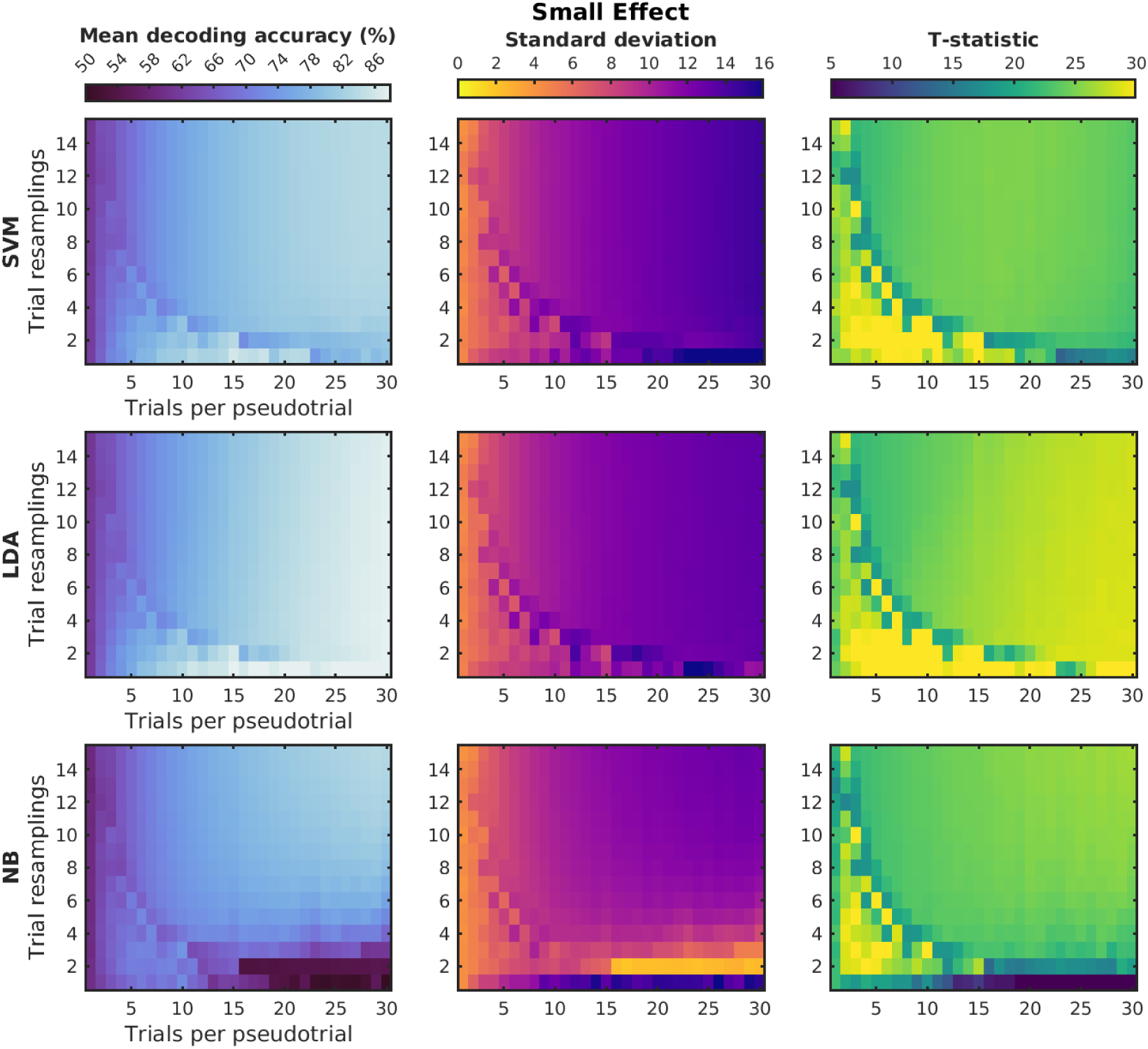
The influence of averaging and resampling using a ‘leave-n-pseudotrials-out’ decoding approach for simulated data with a small underlying effect. Rows correspond to results from the three classifiers tested (SVM = support vector machine, LDA = linear discriminant analysis, NB = Naïve Bayes). Here the number of cross-validation folds was determined by the number of trials per pseudotrial (the more trial averaging and resampling, the fewer possible folds), with trials randomly allocated across 100 iterations of pseudotrial creation. For the results plotted here, we simulated data from 100 subjects with 90 trials per condition and a large effect, or class distance of 0.2 (see main Figure 4 for a small effect, or class distance of 0.1).

**Supplementary Figure 5.**
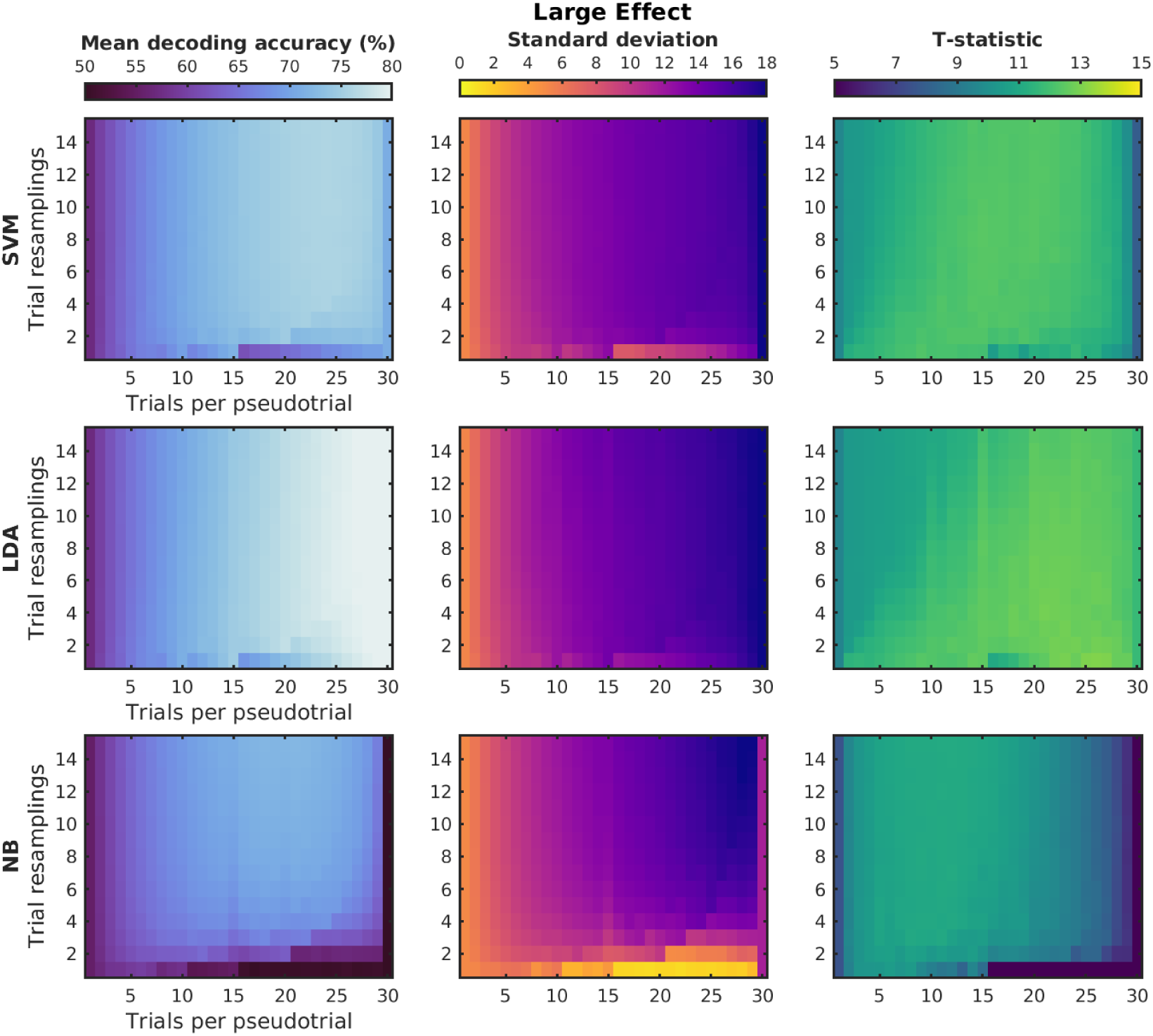
The influence of both averaging and resampling with fewer subjects (n=50) for simulated data with a large underlying effect. Rows correspond to results from the three classifiers tested (SVM = support vector machine, LDA = linear discriminant analysis, NB = Naïve Bayes). Pseudotrials were created separately within 3 allocated ‘blocks’ of trials, facilitating a 3-fold cross-validation approach, with trials randomly allocated across 100 iterations of pseudotrial creation.

For the results plotted here, we simulated data from 50 subjects with 90 trials per condition and a large effect (class distance of 0.2).

### S5. Influence of a smaller number of features

Here we examined the influence of reducing the number of simulated features from 700 to 6, which was not reported in the main text. The overall performance of the classifiers was reduced, as well as the influence of the parameters on t-statistics. However, using around a third of the original trials per pseudotrial and a resampling of 2 would still be a reasonable choice to optimise classification performance.

**Supplementary Figure 6.**
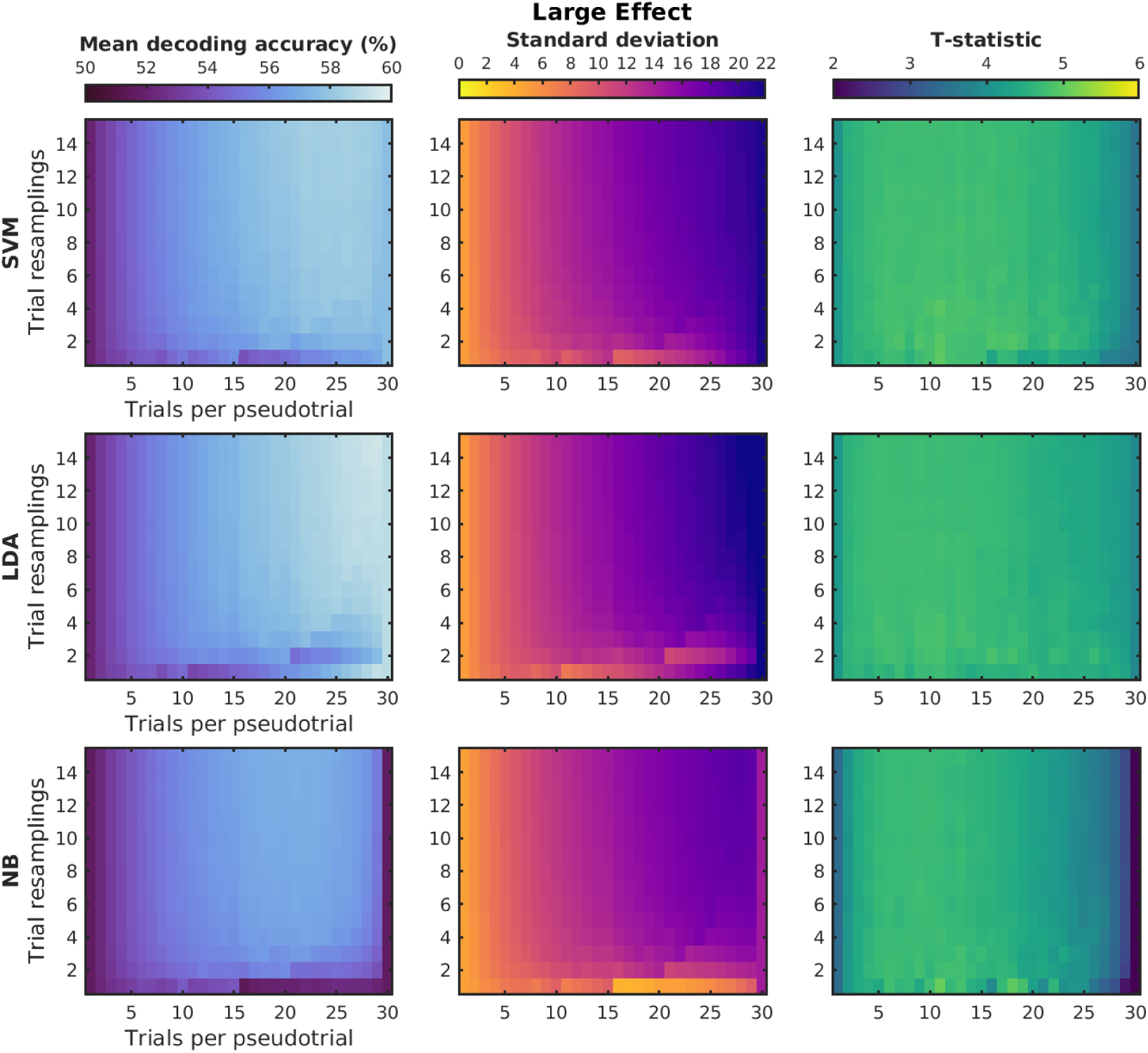
The influence of both averaging and resampling with a smaller number of features (data size ‘small’ = 6 features) for simulated data with a large underlying effect. Rows correspond to results from the three classifiers tested (SVM = support vector machine, LDA = linear discriminant analysis, NB = Naïve Bayes). Pseudotrials were created separately within 3 allocated ‘blocks’ of trials, facilitating a 3-fold cross-validation approach, with trials randomly allocated across 100 iterations of pseudotrial creation. For the results plotted here, we simulated data from 100 subjects with 90 trials per condition and a large effect (class distance of 0.2).

### S6. Influence of increasing the number of cross-validation folds (10-fold)

Here we examined the influence of increasing the number of cross-validation folds from 3 to 10. As we simulated 90 trials per condition, there was a maximum of 9 original trials per pseudotrial. As in the 3-chunk version (**main** Figure 3), using around a third of the original trials per chunk to create each pseudotrial (i.e., 3 trials) was sufficient to aid decoding. However, even for the Naïve Bayes classifier there was little to no benefit of resampling.

**Supplementary Figure 7.**
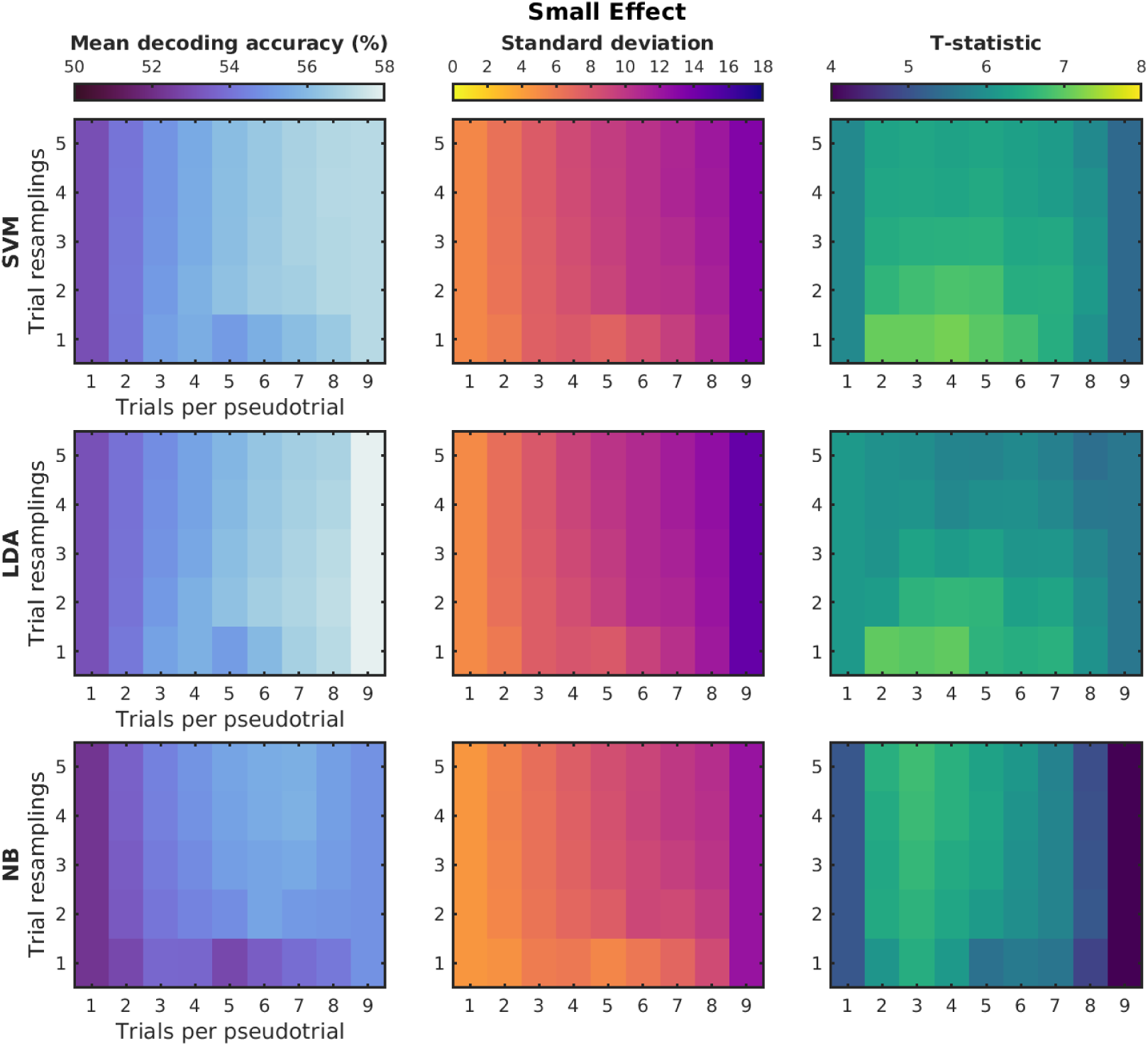
The influence of averaging and resampling using a higher number of cross-validation folds for simulated data with a small underlying effect. Rows correspond to results from the three classifiers tested (SVM = support vector machine, LDA = linear discriminant analysis, NB = Naïve Bayes). Pseudotrials were created separately within 10 allocated ‘blocks’ of 9 trials each, facilitating a 10-fold cross-validation approach, with trials randomly allocated across 100 iterations of pseudotrial creation. For the results plotted here, we simulated data from 100 subjects with 90 trials per condition and a small effect (class distance of 0.1).

### S6. Influence of fewer iterations of random trial allocation

For the results reported in the main text, we ran 100 iterations of the pseudotrial procedure and averaged the 100 resulting classification accuracies. This was to ensure that the results were not dependent on the specific division of trails into pseudotrials, for each ‘subject’ and parameter set. Supplementary Figure 8 demonstrates the pattern of results achieved with one, five, 50, and 100 iterations of random trial allocation. Without this iteration procedure (when the number of iterations = 1), trial averaging quickly increases the between-subject variance, meaning that little or no benefit is derived from the pseudotrial procedure. In fact, the pseudotrial procedure now becomes detrimental in most cases, especially for the NB classifier and for high numbers of trials/pseudotrial for SVM. Thus, including multiple iterations appears to be crucial for deriving benefit from the pseudotrial procedure.

**Supplementary Figure 8.**
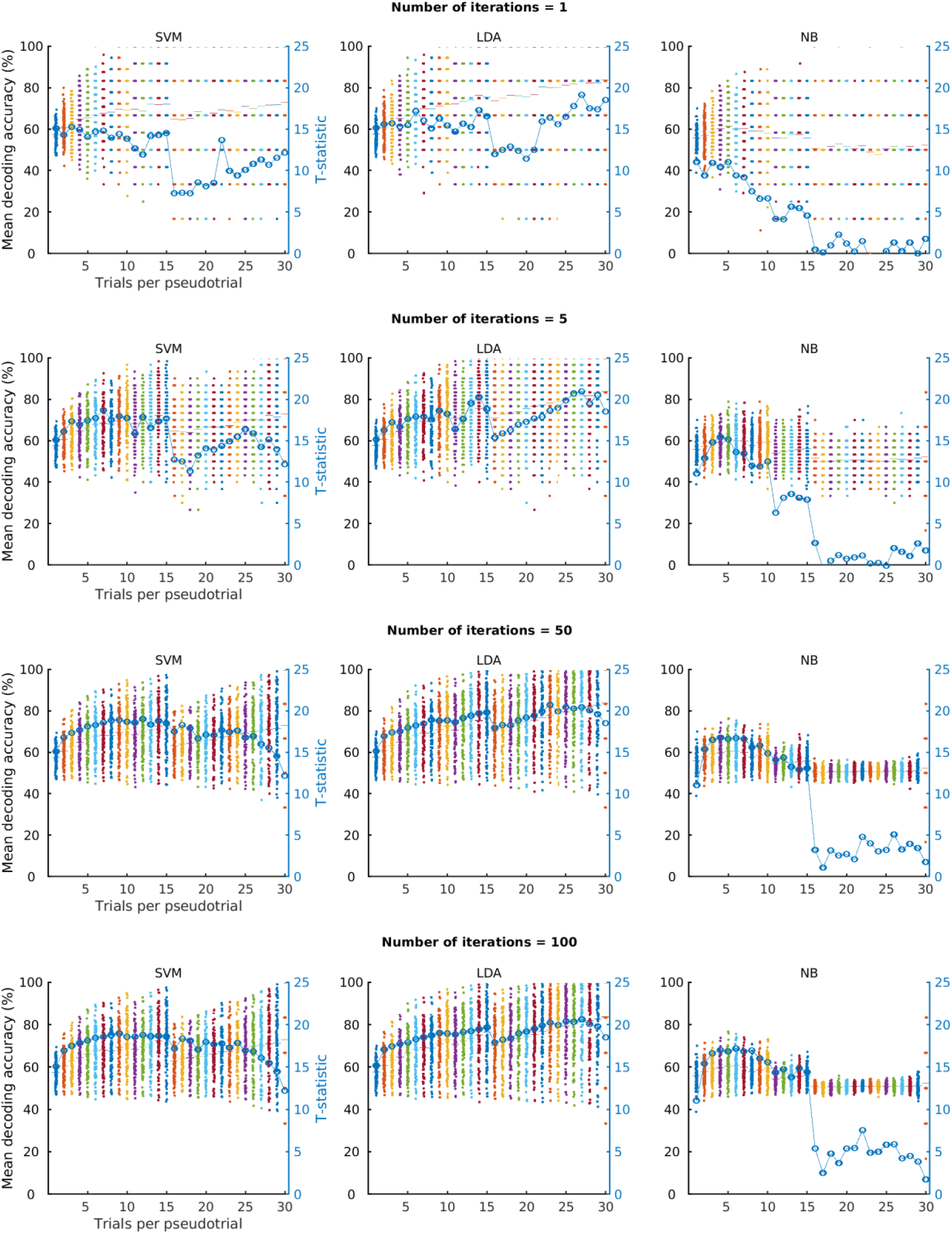
The influence of fewer iterations of random trial allocation for simulated data with a large underlying effect. For the results plotted here, we simulated data from 100 subjects with 90 trials per condition and a large effect (class distance of 0.2). Rows correspond to the number of iterations of random trial allocation that was used to create pseudotrials. Columns correspond to results from the three classifiers tested (SVM = support vector machine, LDA = linear discriminant analysis, NB = Naïve Bayes). Pseudotrials were created separately within 3 allocated ‘blocks’ of trials, facilitating a 3-fold cross-validation approach, with trials randomly allocated.

## Acknowledgements

This work was funded by the MRC Intramural funding, SUAG/093 G116768, and ARC Discovery Project, DP170101840. TG is supported by ARC fellowship DE230100380. For the purpose of open access, the author has applied a Creative Commons Attribution (CC BY) licence to any Author Accepted Manuscript version arising from this submission. Thank you to Dr Arran Reader for their comments on an earlier version of the manuscript.

## Data and Code Availability

Analysis scripts can be found at https://osf.io/hjf75/.

## Author Contributions

**CS:** methodology, software, formal analysis, writing – original draft, visualisation. **TG:** conceptualisation, methodology, software, writing – editing and reviewing, visualisation. **AW:** conceptualisation, methodology, software, writing – editing and reviewing, visualisation, supervision, funding acquisition.

## Declaration of Competing Interests

Nothing to declare.

## Ethics

No ethical approval required for simulated data.

## Notes

https://osf.io/hjf75/

### Competing Interest Statement

The authors have declared no competing interest.

### Summary of Updates

Improved introduction, clearer descriptions and figures.

